# CiiiDER: a new tool for predicting and analysing transcription factor binding sites

**DOI:** 10.1101/599621

**Authors:** Linden J. Gearing, Helen E. Cumming, Ross Chapman, Alexander M. Finkel, Isaac B. Woodhouse, Kevin Luu, Jodee A. Gould, Samuel C. Forster, Paul J. Hertzog

**Author notes:** These authors contributed equally to this work (Linden J. Gearing, Helen E. Cumming). These authors also contributed equally to this work (Samuel C. Forster, Paul J. Hertzog).

## Abstract

The availability of large amounts of high-throughput genomic, transcriptomic and epigenomic data has provided opportunity to understand regulation of the cellular transcriptome with an unprecedented level of detail. As a result, research has advanced from identifying gene expression patterns associated with particular conditions to elucidating signalling pathways that regulate expression. There are over 1,000 transcription factors (TFs) in vertebrates that play a role in this regulation. Determining which of these are likely to be controlling a set of genes can be assisted by computational prediction, utilising experimentally verified binding site motifs.

Here we present CiiiDER, an integrated computational toolkit for transcription factor binding analysis, written in the Java programming language, to make it independent of computer operating system. It is operated through an intuitive graphical user interface with interactive, high-quality visual outputs, making it accessible to all researchers. CiiiDER predicts transcription factor binding sites (TFBSs) across regulatory regions of interest, such as promoters and enhancers derived from any species. It can perform an enrichment analysis to identify TFs that are significantly over- or under-represented in comparison to a bespoke background set and thereby elucidate pathways regulating sets of genes of pathophysiological importance.

CiiiDER is available from www.ciiider.org.

## Introduction

Contemporary transcriptomic technologies such as microarrays and RNA-sequencing provide reliable methods to identify genes differentially expressed across cell types, tissues or in response to different stimuli. These methods reveal many co-expressed genes or gene networks that are together predicted to determine the observed biological responses.

Transcription factors (TFs) bind to specific DNA sequences (transcription factor binding sites; TFBSs) within promoter and enhancer regions of genomic DNA and either activate or repress gene expression. These interactions can be determined experimentally, for example using chromatin immunoprecipitation (ChIP) techniques, and are typically represented as position frequency matrices (PFMs). Curated databases of PFMs, applicable to a wide range of species, include the commercial TRANSFAC database [1] and the open-access JASPAR database [2, 3]. While using PFMs alone can predict TFBSs within regulatory sequences, in eukaryotic organisms, this typically results in high false positive prediction rates, since predicted sites may not be accessible to the transcriptional machinery due to chromatin structure or the epigenetic landscape.

An enrichment analysis, which compares the distribution of TFBSs predicted in a set of regulatory DNA regions to the distribution in a set of background sequences, can be utilised to more accurately identify true TFBSs. With the appropriate choice of background, it is possible to identify TFBSs that are statistically over- or under-represented. The TFs with over-represented TFBSs in a set of co-expressed genes are more likely to be involved in regulating the expression of these genes.

While there are existing and publicly available programs that can perform enrichment analyses, these tools are typically web-based (e.g. Pscan [4], MotifViz [5] and oPOSSUM [6]) or restricted to command line use (e.g. Clover [7], the MEME suite [8] and HOMER [9]). Online tools can be convenient, but it is beneficial to be have the security of a downloadable program, which can be run on a local computer and to have saved projects that can easily be revisited. Downloadable applications for TFBS prediction that are run through the command line often require additional tools to be installed or lack effective visualisations, providing static or text-based results, which limits their utility for a wide audience.

There is a need for a downloadable program, independent of computer operating system, that provides the required flexibility to perform accurate, integrated, customisable analysis and data exploration. To address this, we developed CiiiDER, a user-friendly analysis toolkit for predicting and analysing putative TFBSs within regulatory regions. CiiiDER is operated through an intuitive graphical user interface (GUI), which is designed with ease of use in mind. Established algorithms [10] have been implemented to map potential TFBSs within a set of regulatory sequences. TF enrichment analysis can be used to enable identification of key TFs that are statistically enriched and are therefore more likely to be biologically relevant.

## Methods

### Workflow and algorithms

The CiiiDER workflow (Fig 1) can accept query and background regions in a variety of input formats. It consists of two main analyses, with results presented in an interactive format.

**Fig 1.**
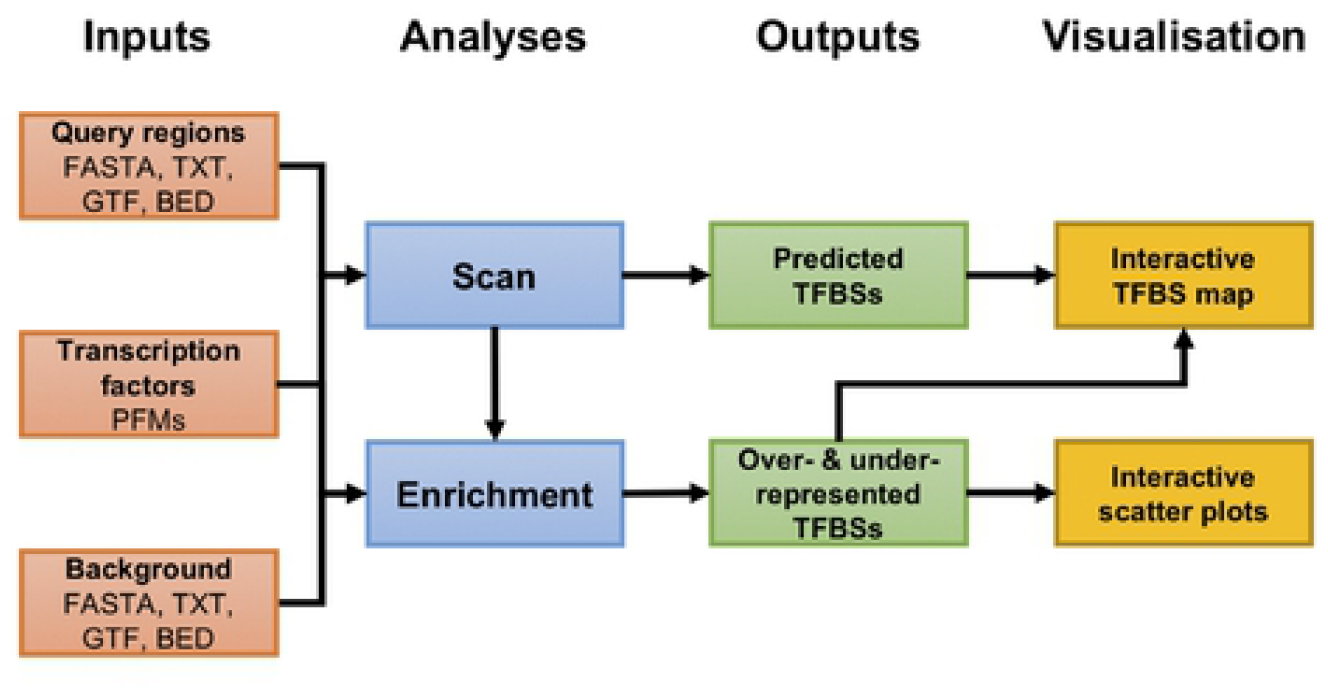
CiiiDER workflow. A typical analysis involves submitting gene sets for scanning against known TF models, followed by the identification of sites that are statistically enriched relative to a submitted background gene list.

#### Inputs

CiiiDER has been designed to ensure that data input is easy and that a wide variety of formats are accepted. All data input and parameter selection is facilitated though simple interfaces, with pop-up information boxes available to help users make the appropriate selections. Data can be loaded from file or by pasting the sequence information directly into a box in the GUI.

CiiiDER will read sequences directly from data entered in FASTA format. Genomic location formats (GFF, GTF or BED) require an associated genome file for the relevant species to obtain DNA sequences. These formats can be used to analyse any regulatory region of interest including promoters, enhancers and untranslated regions (UTRs).

As promoters tend to be the regulatory regions of interest for the majority of users, CiiiDER will automatically extract promoter sequences from a genome file, given a list of gene symbols, Ensembl gene IDs or Ensembl transcript IDs. This requires an additional annotation file, denoted the gene look-up manager (GLM), which contains the location of the transcription start sites (TSSs) for each gene and transcript. The user can specify the size of the sequence relative to the TSS, with the default set at −1500 bases upstream to +500 bases downstream of the TSS. CiiiDER can be downloaded with human GRCh38 and mouse GRCm38 genomes and GLM files. For alternative genomes, the user will need to provide the appropriate genome and an Ensembl GTF file, from which CiiiDER can automatically generate the necessary GLM file.

#### Scan

During the Scan stage, CiiiDER uses an implementation of the MATCH algorithm [10] to predict potential TFBSs in regions of interest. This approach is compatible with PFMs in JASPAR [11] or TRANSFAC [1] format. The mapping of each TFBS is performed with a user-specified deficit that determines the stringency of the scan. The deficit is the difference between the MATCH score of a TFBS and the maximum possible score, which is 1. The default deficit is 0.15, which means the scan will accept any TFBSs that have MATCH scores of 0.85 or above (the same cut-off is applied to both the core and matrix scores from the MATCH algorithm).

#### Enrichment

The Enrichment stage identifies those TFBSs that are significantly over- or under-represented in query regions compared to relevant, user-specified background regions. This analysis scans the background sequences for TFBSs using the same criteria as used for the query sequences. Over- and under-represented TFs are determined by comparing the numbers of sequences with predicted TFBSs to the number of those without, using a Fisher’s exact test; alternatively, the distributions of the number of sites per sequence in the query and background sets can be compared using a Mann-Whitney U test.

For the enrichment plots, if a given transcription factor has binding sites in *n_S_* out of *N_S_* search regions and *n_B_* out of *N_B_* background regions, then:

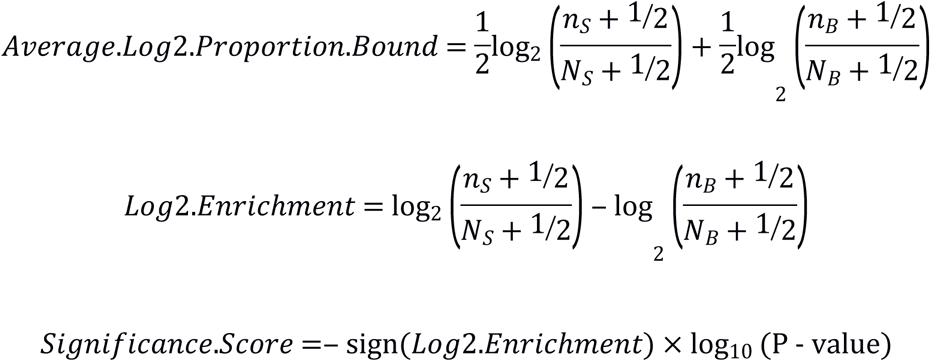

#### Outputs and Visualisation

CiiiDER produces clear graphical displays to help interpret the results of TFBS prediction. Putative TFBSs are displayed in an interactive map (**Error! Reference source not found.**). By default, ten TFs are shown, but the user can choose to add or remove TFs from the image. There are also options to filter the TFs displayed according to the scan stringency or enrichment *P*-value, for intuitive exploration of the data.

Promoters or other regulatory regions can be re-arranged or removed and the colour of each TF can be customised for the production of figures. The enrichment analysis produces an additional interactive plot that displays the fold enrichment, average abundance and *P*-value associated with all TFs (Fig 3). The images created can be saved as publication-quality files and the binding site data and enrichment statistics can be saved as text files for subsequent analysis using additional tools.

The entire project can be saved for further analysis and interpretation.

**Fig 2.**
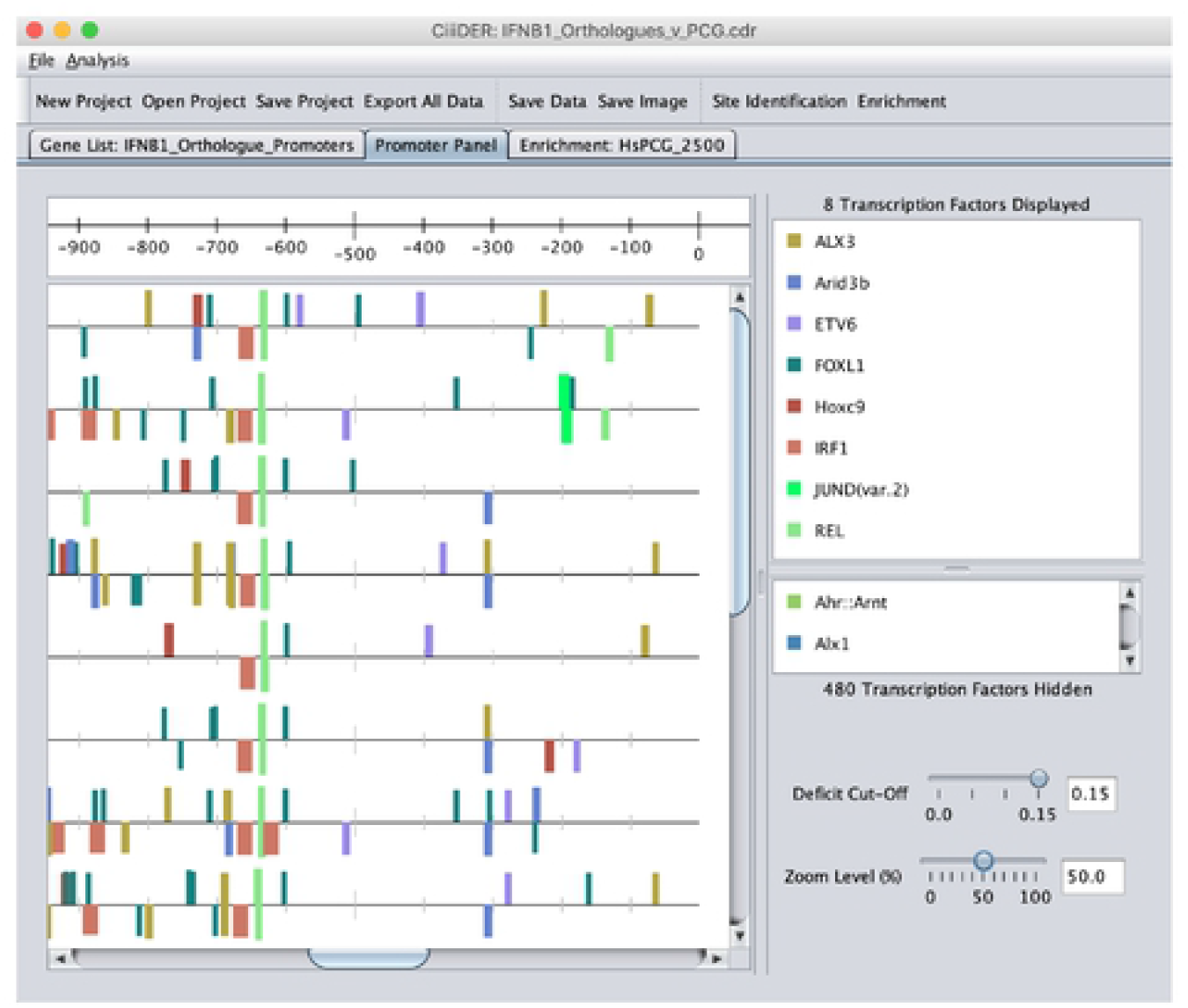
CiiiDER interactive site map. The scan and enrichment algorithms produce a graphical display of the TFBS locations on the sequences. There are many options to edit the images, including adjusting the deficit and *P*-value thresholds for displaying TFBSs, selecting or removing TFs to be viewed, editing the colour scheme for TFs and rearranging the order of the sequences.

**Fig 3.**
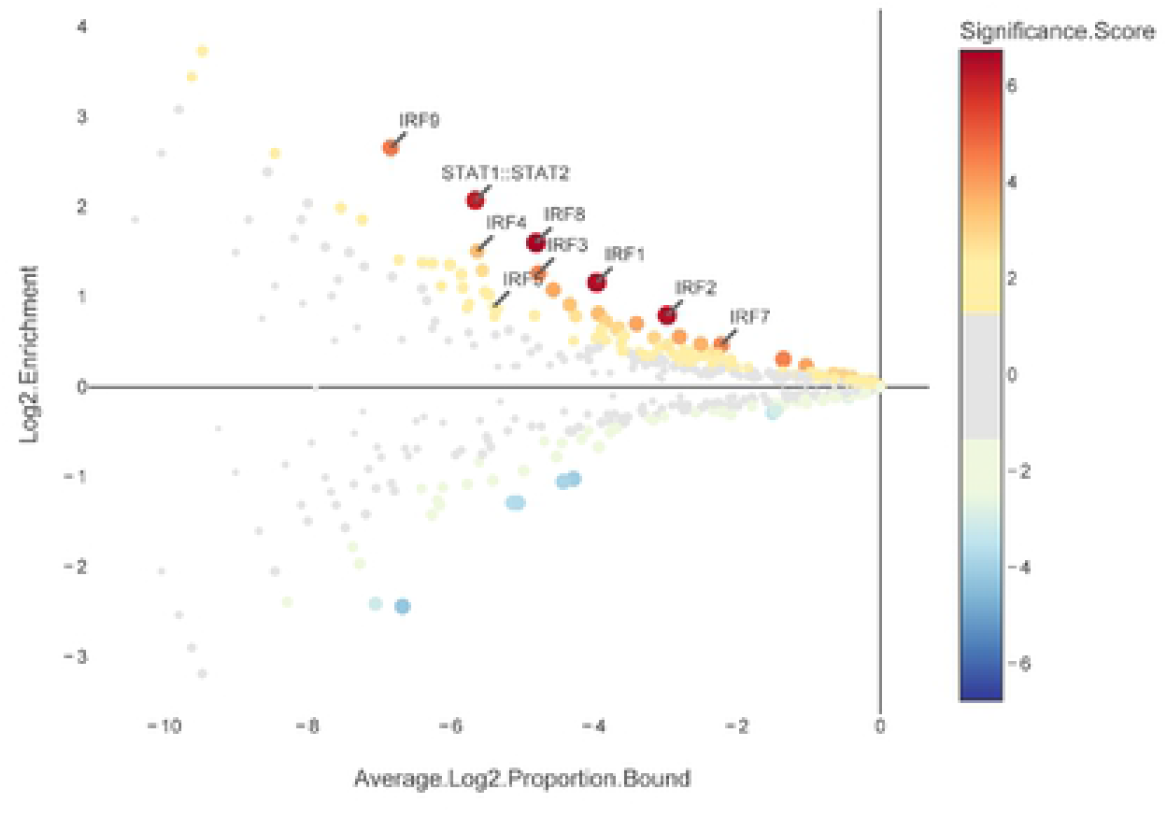
CiiiDER enrichment results for the breast cancer metastasis dataset. The data are derived from the proportion of regions bound for each TF, which is the number of bound regions divided by the total number of regions. The plot shows the enrichment (ratio of proportion bound) and average log proportion bound. Size and colour show ∓log_10_(*P*-value); it is greater than zero if the TF is over-represented and less than zero if under-represented.

#### Implementation

CiiiDER has been implemented in Java for platform independence with the GUI utilising the Swing libraries to deploy a simple, intuitive interface. The multi-threading capabilities of Java are used to take advantage of all available computer processors, significantly improving analysis speeds (S1 Fig); alternatively, CiiiDER can also be restricted to use only a certain number of processors. Enrichment of transcription factors is also displayed using interactive HTML plots generated using the Plotly JavaScript library [12].

CiiiDER is available for download as a JAR file with supporting files; other software dependencies do not need to be installed. Two PFM libraries are supplied with the software: the JASPAR 2018 CORE non-redundant vertebrate matrices [2] and matrices from Jolma *et al.*, a large experimental dataset [13]. Genomes and associated GLM files are also available for extracting promoter sequences using gene names or Ensembl IDs.

CiiiDER is distributed under the GNU GPLv3 licence. The program and documentation are available from www.ciiider.org and the source code is available at https://gitlab.erc.monash.edu.au/ciiid/ciiider.

### Experimental methods

CiiiDER analyses were performed using the Ensembl 89 or Ensembl 94 annotations of the human GRCh38 and mouse GRCm38 genomes, respectively, with the 2011 version of the TRANSFAC non-redundant vertebrate database [1] or the 2018 JASPAR core non-redundant vertebrate matrices [2]. All promoter regions were defined as spanning −1500 bases to +500 bases relative to the transcription start site.

#### ChIP-seq scan

Experimentally verified transcription factor binding site regions were obtained from publicly available ChIP-seq experiments: a CTCF dataset (ENCSR000DLG) from ENCODE [14] and STAT3 dataset (GSM288353 [15]) from GEO [16]. These were selected because they were in narrow peak format with peak max values, to give the highest probability of focusing on the true binding site, and because matching high-quality TRANSFAC TF models were available.

Sequences corresponding to 50 bases either side of the maximum signal of each ChIP-seq peak were obtained. This length was chosen to allow sufficient sequence to identify TFBSs, while minimising extraneous sequence. Backgrounds were produced by using 101 base genomic sequences 10,000 bases away from the peak, ensuring that none of the background sequences overlapped with surrounding peaks. CiiiDER scans were performed using deficits of 0.2. TRANSFAC scans were performed with equivalent core and matrix similarity score cut-offs of 0.8. Clover analyses were performed with default values. pROC [17] was used to generate ROC curves and associated area under curve (AUC) values for each program.

#### Microarray enrichment analysis

Robust multiarray averaging (RMA)-normalised microarray data from Bidwell *et al.* (GSE37975) [18] were downloaded using GEOquery [19] and processed through the limma package [20] to identify differentially expressed transcripts between the primary and metastatic tumour samples, with log_2_ fold change > 1 or < −1 and a *P*-value < 0.05 (adjusted for false discovery rate using the Benjamini-Hochberg correction).

The query gene list consisted of significantly down-regulated genes that were defined as interferon-inducible in mouse using the Interferome v2.0 [21] (up-regulated at least two-fold by interferon treatment). The background gene list was derived from transcripts with an absolute log_2_ fold change < 0.05 and average expression greater than the first quartile. CiiiDER analyses were performed on the promoter sequences using deficits of 0.15 with JASPAR TF models.

#### Phylogenetic scan

The IFNβ promoter sequence was obtained from 16 species of placental mammals, using Ensembl 94 annotations. A background containing the promoter sequences of 2,463 human protein-coding genes was used for enrichment. CiiiDER analysis was performed using JASPAR TF motifs with deficits of 0.15.

## Results and discussion

### Comparing CiiiDER to other software with ChIP-seq data

In order to demonstrate the utility of CiiiDER and that it correctly implements the MATCH algorithm, we compared CiiiDER against other downloadable TF site prediction software, TRANSFAC (which also uses the MATCH algorithm), as well as Clover. As examples, we obtained ChIP-seq data for CTCF and STAT3 and compared the performances of the programs to detect true positive TFBSs using ROC curves (**Error! Reference source not found.**.). The Area Under Curve (AUC) was calculated for each program and CiiiDER showed the same ability to predict true positive sites as TRANSFAC, demonstrating that the CiiiDER implementation of the MATCH TFBS prediction algorithm functions as expected.

### Applications of CiiiDER

#### Identification of enriched TFs in co-expressed gene sets

Transcriptomic experiments such as microarray and RNA-sequencing are an excellent source of co-regulated genes for CiiiDER analyses. In order to perform an enrichment analysis for a gene set of interest, it is important to choose an appropriate background. Comparing an experimentally derived, co-expressed gene list to a genome-wide background may lead to the enrichment of some TFs that are not specifically related to the experiment. For example, if the query were promoters of genes showing a significant change in expression following a particular treatment, then an appropriate background might be promoters of genes that were expressed in the appropriate cell or tissue type, but showed no significant response to the stimulation.

If the experiment were comparing two cell types, the query could be genes expressed highly in one cell type and the background would then be genes expressed highly in the other. Over represented TFs would be predicted to be important in regulating transcription in the first cell type, whereas under-represented TFs could be important in transcriptional regulation in the second cell type.

The ability for CiiiDER to predict key regulatory TFs was demonstrated by reanalysing a published study of the regulation of the immune system in breast cancer [18]. Bidwell *et al.* compared the gene expression in primary and metastatic tumour cells in a mouse model of spontaneous bone metastasis. A set of approximately 3,000 genes were down-regulated in the metastasised cells relative to the primary tumour, 540 of which were determined to be interferon-regulated genes (IRGs) from the Interferome v1.0 database [22]. In that study, Clover was used to show that these genes were enriched for Irf7 binding sites. The role of Irf7 was confirmed by showing an increase in interferon signalling and a reduction of tumour metastases following restoration of Irf7 expression in the tumour cells in the bone metastasis model.

We reanalysed the normalised expression data from this experiment to create a list of IRGs down-regulated in metastases using the updated Interferome v2.0 [21]. A CiiiDER enrichment analysis was performed on the promoters of these IRGs using a background gene set of expressed, unchanged genes (Fig 3). This showed that CiiiDER was able to identify Irf7 as a key TF, potentially regulating the expression of immune system genes within the breast cancer tumour (*P* = 3.46E-05), in agreement with the published prediction and experimental validation.

In this example, many other IRF-family TFs were also significantly over-represented (e.g. IRF8, *P* = 1.76E-07). It is often difficult to accurately distinguish between TFBSs of TFs belonging to a family, since their binding site preferences can be very similar. In this case, cross-referencing with the published expression data revealed that Irf7 was the most significantly suppressed IRF-family TF in metastases, which added supporting evidence to its role.

#### Identifying phylogenetically conserved TFBSs

Since gene orthologues often retain similar functions throughout evolution and maintain a similar method of regulation [23], CiiiDER could potentially be used to examine phylogenetic conservation, through prediction of enriched TFs, and by creating visualisations to help distinguish patterns in TFBSs. To test the capacity of CiiiDER to identify evolutionary conserved regulatory elements, we selected the interferon β (IFNβ) promoter, the transcriptional regulation of which has been very well characterised [24]: in brief, IRF-family TFs, NF-κB and AP-1 (which is comprised of ATF2 and JUN) together allow remodelling of the local chromatin structure to promote gene transcription.

Initially, the scan method was used to identify TFBSs in the IFNβ promoters from placental mammal species detailed in Fig 4. This identified a great number of potential TFBSs for hundreds of TFs (see Fig 2), many of which are likely to be false positives, which makes it difficult to identify likely candidate transcriptional regulators. An enrichment analysis was then performed to compare the TFBSs in the IFNβ promoters against a background of human protein coding genes. The top ten over-represented TFs that occurred in at least half of the promoters were selected for display (Fig 4). Spatially conserved TFBSs are immediately apparent, particularly those in a cluster within 200 bases of the TSSs. This includes TFBSs for the best characterised IFNβ regulators—NF-κB components RELA and REL (*P* = 2.38E-19 and 1.10E-10) and IRF-family TFs IRF1 and IRF2 (*P* = 9.54E-14 and 4.46E-10). These are the most significant TFs that are predicted in all promoters, whereas other top significant TFs do not show the same consistent pattern.

**Fig 4.**
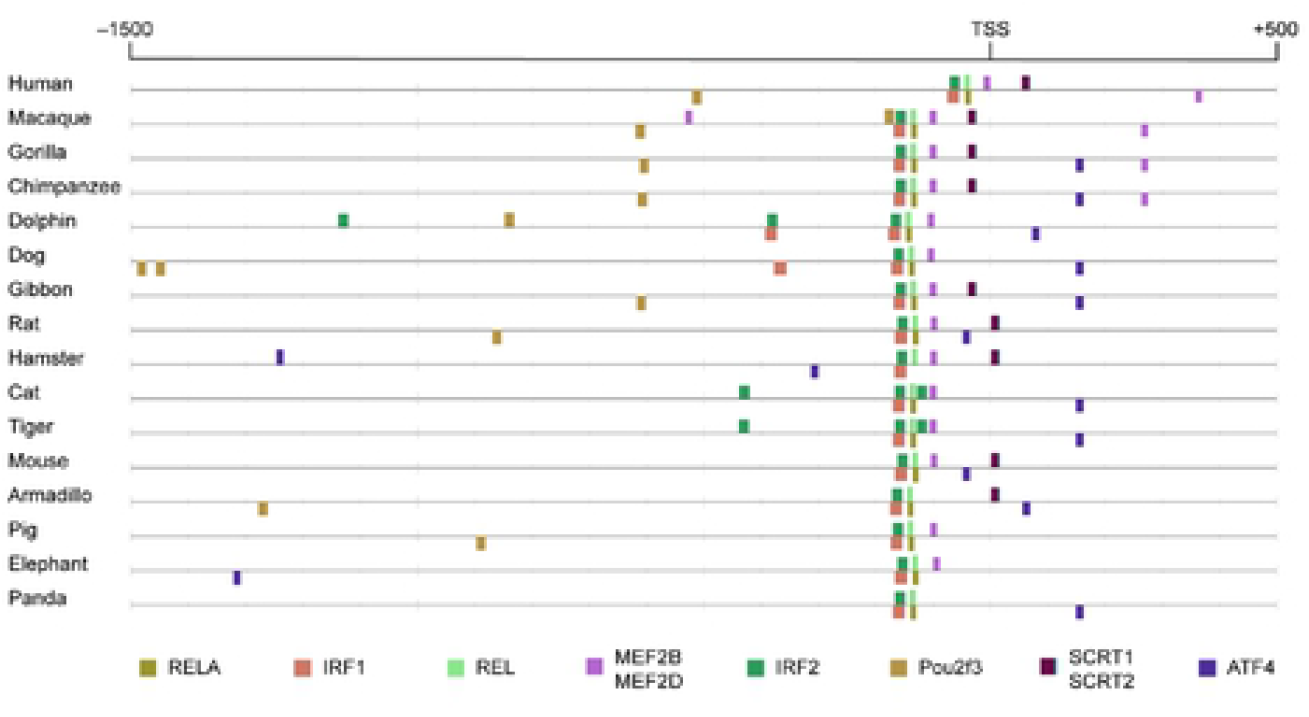
Phylogenetic conservation of TF binding sites in the IFNβ promoter. The results of the enrichment algorithm, displaying the ten most significantly enriched TFs present in at least half of the promoters.

The combination of enrichment analysis and effective visualisation can allow rapid identification of TFBSs that are phylogenetically and spatially conserved. This gives greater support when choosing candidate TFs that are most likely to be involved in regulatory elements.

### Further analyses

CiiiDER can also be used to search for TFBSs associated with regions of the genome marked with epigenetic modifications obtained from ChIP-seq data or open chromatin regions derived from ATAC-seq data. For example, we have published using CiiiDER to examine transcriptional enhancers (marked by histone H3 lysine 4 mono-, di- and tri-methylation) in effector and memory T-cells, compared to those common between effector, memory and naïve T-cells, to show an enrichment of BATF, JUN and FOS motifs, among others [25].

The power of CiiiDER analyses can be increased by linking the results to other data. As with the breast cancer example, it is worth considering all members of a TF family when choosing TFs for further validation. TFBS enrichment results may be assessed in the context of gene expression data to determine which TFs are detectable or have altered expression levels in the experimental system of interest.

## Conclusion

CiiiDER is an intuitive new tool for analysing TFBSs in regulatory regions of interest. It can efficiently scan sequences for potential TFBSs and identify TFBSs that are statistically under- or over-represented. It is user-friendly and produces quality visual outputs to assist researchers to uncover signalling pathways and their controlling TFs in a wide variety of biological contexts. The program, user manual and example data are available at www.ciiider.org.

## Acknowledgements

The authors would like to thank Gordon Smyth and Jonathan Keith for statistical advice and Michael Gantier, Mirana Ramialison, Christine Wells and Jesse Balic for evaluating the software. The authors would like to acknowledge the support and contributions of the Bio21 Undergraduate Research Program, the Victorian Life Sciences Computation Initiative, Monash e-Research and Philip Chan for support in this project. Research at Hudson Institute of Medical Research is supported by the Victorian Government’s Operational Infrastructure Support Program.

## Supporting information captions

**S1 Fig.**
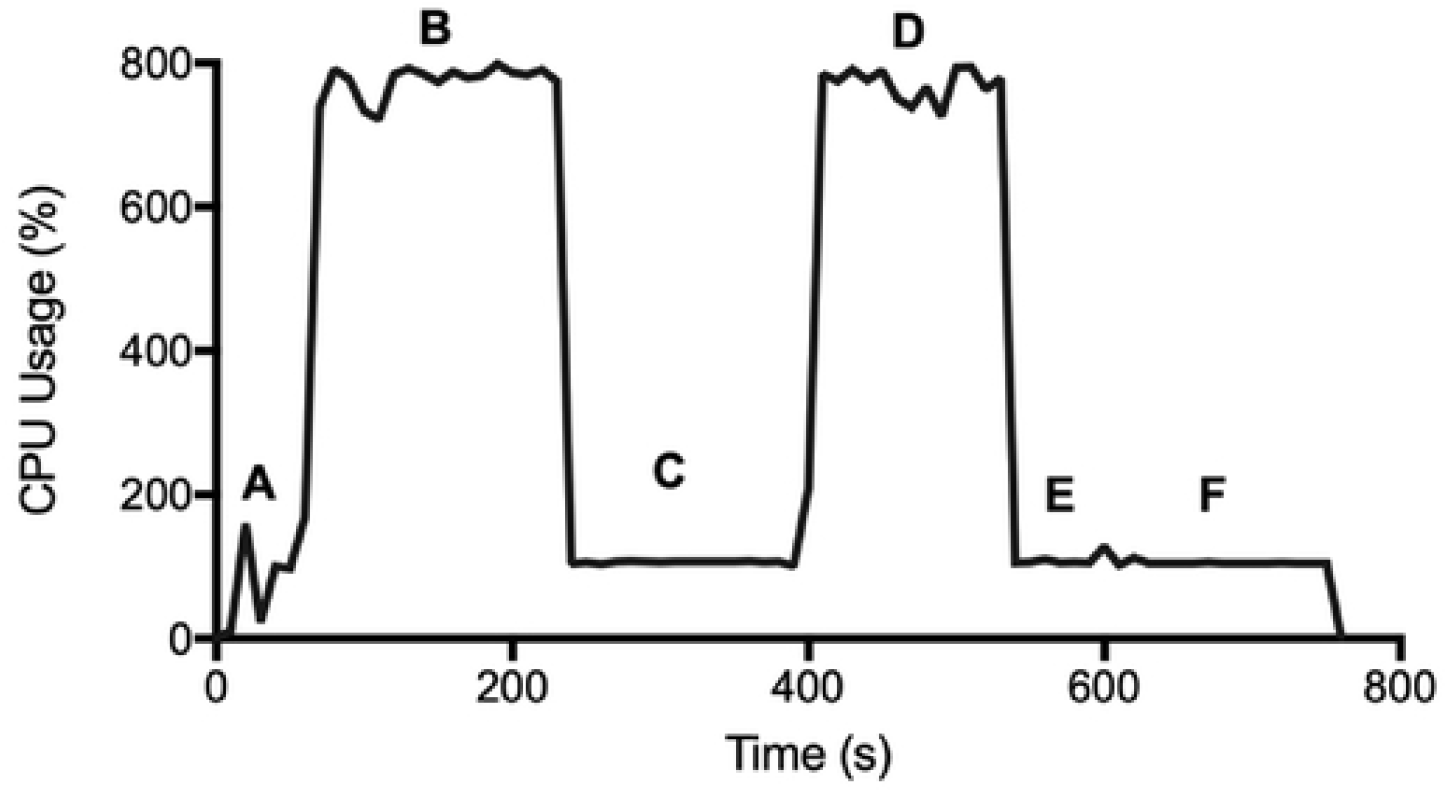
CiiiDER analysis time. Example plot of analysis time and CPU usage of CiiiDER when performing site identification and enrichment using the Irf7 breast cancer gene set. Gene sets were loaded into the GUI and promoters were obtained (A), TFBSs were predicted across the query promoters (B) and collated (C), background sites were predicted (D), the enrichment calculation was performed (E) and the final graphical outputs were created (F). The site prediction steps take advantage of multiple computer processors. The maximum memory usage was 4.53 GB. Measurements were made on an iMac with four i7 4.0 GHz processors and 32 GB RAM.

**S2 Fig.**
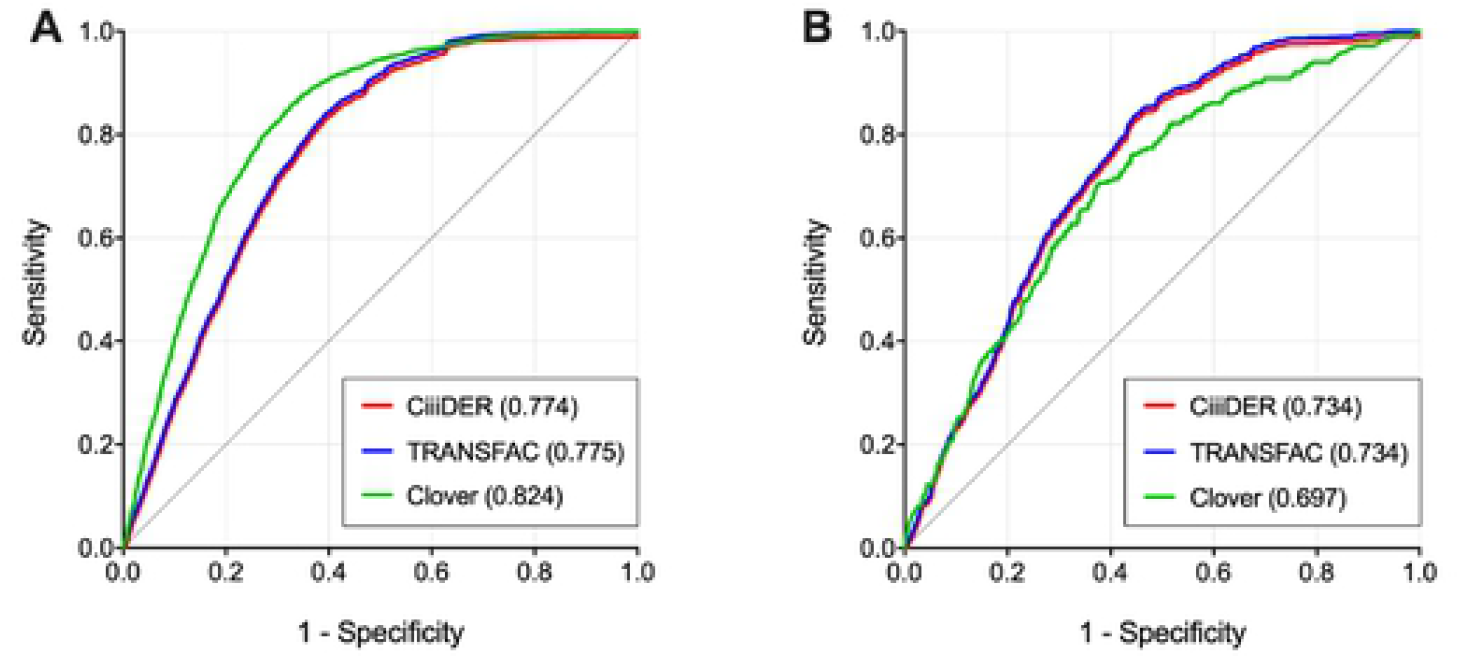
CiiiDER scan algorithm. The accuracy of CiiiDER was compared with Clover and TRANSFAC software using ROC curves for (A) CTCF and (B) STAT3. The curves represent the ratio of true binding sites predicted against the number of false binding sites predicted. The locations of true binding sites have been validated previously using ChIP-seq experiments. Note that, due to almost complete overlap with the TRANSFAC curves, the CiiiDER curves for both CTCF and STAT3 were shifted down by −0.01.

